# Independent evolution of cutaneous lymphoma subclones in different microenvironments of the skin

**DOI:** 10.1101/829671

**Authors:** Aishwarya Iyer, Dylan Hennessey, Sandra O’Keefe, Jordan Patterson, Weiwei Wang, Gane Ka-Shu Wong, Robert Gniadecki

## Abstract

Mycosis fungoides (MF) is the most common, yet incurable, cutaneous T-cell lymphoma. We have recently shown that the disease is initiated by hematogenous seeding the skin with clonotypically diverse neoplastic T-cells which proliferate accumulating numerous mutations and produce lesions of high intratumoral heterogeneity (ITH). A characteristic but a poorly studied feature of MF is epidermotropism, the tendency to infiltrate skin epithelial layer (epidermis) in addition to the vascularized dermis. By sequencing the exomes of the microdissected clusters of lymphoma cells from the epidermis and the dermis, we found that those microenvironments harbored different malignant clonotypes and exhibited different patterns of driver gene mutation. Phylogenetic relationships between cancer subclones witnessed to the independent mutational evolution in the epidermis and dermis. Thus, the invasion of MF to different skin layers does not occur by gradual infiltration of the expanding tumor mass, but is caused by separate seeding processes with different malignant clones that develop independently of one another via a neutral, branched evolution. In conclusion, tissue microenvironments shape the subclonal architecture in MF leading to “ecological heterogeneity” which contributes to the total ITH. Since ITH adversely affects cancer prognosis, targeting the microenvironment may present therapeutic opportunities in MF and other cancers.

## Introduction

Mycosis fungoides (MF) is one of the most common diseases in the realm of extranodal T-cell lymphomas (1). It is a skin-tropic lymphoid neoplasm that initially presents as scaly, erythematous patches and plaques, which may progress to tumours and disseminate to lymph nodes and other organs (2–4).

ITH has recently emerged as an important characteristic of solid and hematopoietic malignancies (5). Although mutations in few driver genes may be sufficient to initiate tumorigenesis, it is now evident that the progression depends on the accumulation of multiple mutations to promote expansion and invasion of the primary niche and surrounding tissues (6,7). Mutations occur randomly in malignant cells within the tumour, leading to the emergence of multiple subclones. ITH allows cancer to withstand selection pressure from the microenvironment and therapies by promoting the expansion of subclones harboring mutations advantageous to these cells (6).

The generation of ITH is usually viewed as an evolutionary process with a single transformed cell as a starting point. This cell proliferates and branches into phylogenetically related subclones (supplementary **Fig S1**) that infiltrate the tissue. Although this model may be applicable to cancers that grow expansively as single tumours, this is not necessarily true for all malignancies. Many cancers comprise the entire ecosystem of primary and metastatic lesions that are physically separated from each other. It has been shown that in such situations, tumour heterogeneity may be augmented by cross-seeding by circulating, genetically diverse cancer subclones, for example, cancer self-seeding by the cells from the metastatic lesion re-entering the primary tumour (8). We hypothesized that a similar mechanism may operate at the microscopic scale for primary cancers, where different compartments within an organ can be colonized by different cancer subclones. Independent seeding of different microscopic compartments within the same organ would increase the heterogeneity of the entire lesion beyond what would have been possible by a continuous evolution from only one ancestral clone (supplementary **Fig S1**).

MF provides with a convenient model to test this hypothesis. The skin has a simple layered structure comprising ectodermal derived epidermis and the mesodermal dermis. Both layers can be occupied by cancer cells in MF. The histopathology of MF reveals disconnected areas of malignant cell clusters in the dermis and the epidermis. Dermal infiltrate is usually perivascular or diffuse whereas lymphoma foci in the epidermis form well-demarcated clusters of cells known as Pautrier abscesses (**Fig S2**). Pautrier microabscesses are a characteristic feature of MF and are present in approximately 20% of all biopsies (9,10). Unlike the dermal perivascular infiltrates that comprise a significant proportion of reactive cells, Pautrier abscesses are believed to contain predominantly cancer cells with a minor admixture of apoptotic Langerhans cells and eosinophils (11). The initial points of entry of malignant cells are the capillaries in the upper (papillary) dermis. Therefore, Pautrier microabscesses are a manifestation of the infiltrative growth in MF by which “epidermotropic” subclones migrate from the dermis to the epidermis.

We and others have recently studied the heterogeneity of MF on the genomic, transcriptomic and cellular levels (12–17). In contrast to previous views considering MF as a relatively simple, monoclonal lymphoproliferation derived from a mature T-cell, we showed that MF comprises multiple mature T-cell clones which undergo branched evolution producing generations of cancer subclones (16, 17). Interestingly, there seems to be very little competition between different subclones and the disease progression is associated with an increase in subclonal diversity rather than a selection of the fittest subclones. We have therefore asked whether different microcompartments in the skin (epidermis vs dermis) play a role in the generation of ITH in MF. We found that Pautrier microabscesses do not comprise a subpopulation of the dermal malignant cells emigrating to the epidermis, but that they originate independently from distinct seeding event and undergo autonomous evolution, reminiscent of the parapatric type of ecological speciation of the organisms.

## Material and Methods

### Sample collection, cryosectioning, laser capture microdissection (LCM) and sample preparation for whole-exome sequencing (WES)

Samples (4mm punch biopsy and 10ml of blood) were obtained from 7 patients under ethics certificate HREBA.CC-16-0820-REN1 approved by Health Research Ethics Board of Alberta, Cancer Committee. Peripheral blood mononuclear cells (PBMC) were used as normal control except in sample MF18 where the epidermal cells were used as normal control for data analysis. Frozen biopsies were sectioned at 10 µm, transferred on 2 µm PEN membrane slides and stained with hematoxylin and eosin. Clusters of atypical cells representing malignant lymphocytes were microdissected from the dermis and the epidermis under 20x or 40x magnification in Leica DM6000B microscope (Wetzlar, Germany). The microdissected epidermal lymphocytes represented Pautrier microabscesses which could readily be identified based on their enlarged hyperchromatic nuclei, lighter cytoplasm and a cleavage separating them from the surrounding epidermis (supplementary **Fig S2**). Sequencing libraries were prepared with NEBNext® Ultra^TM^ II kit for Illumina (cat# E7645S) (New England Biolabs, MA) and exomes were captured with SSELXT Human All exon V6 +UTR probes (Agilent Technologies, CA). Samples were sequenced on Illumina HiSeq 1500 sequencer or NovaSeq 6000 platform. Detailed protocol for samples processing for storage and sequencing explained in previous methods (15).

### Data analysis

To identify the TCR sequences, the fastq files were analyzed using MiXCR (version 2.10.0) (18). To identify the genomic subclones, the sequenced reads were processed using the GATK4 (version 4.0.10). Somatic variants (SVs) were identified by MuTect2 (version 2.1) (19,20) and Strelka2 (version 2.9.10) (21). Variants filtered as “Pass” from both variant callers were used for downstream analysis. Variant effect predictor (VEP, version 95.2) was used to assign functional significance to the predicted SVs (22). Titan-CNA (version 1.20.1) was used to identify copy number aberrations (CNA) and predict the tumour cell fraction (TCF) (22,23). PhyloWGS (version 1.0-rc2) was used for phylogenetic analysis of the genetic subclones (22–24).

## Results

### Clonotypic diversity of malignant T-cells in epidermal and dermal niches in the skin

Since the epidermis is not vascularized, the intraepidermal neoplastic cells of Pautrier microabscesses must necessarily originate from the cells that initially enter the papillary dermis. Therefore, Pautrier microabscesses are assumed to represent a fraction of the dermal cells that acquired an ability to survive and proliferate in the epidermis. To examine this hypothesis we microdissected atypical cells from both layers (epidermal and dermal) of the skin in 7 MF patients (supplementary **Table S1**) and analysed their clonotypic composition by comparing their T-cell receptor β (TCRβ) repertoires. We used the previously described methodology where the CDR3 sequences of the rearranged *TCRB* genes are detected by bioinformatic analysis of WES data (15). Since *TCRB* locus is only rearranged on one chromosome (allelic exclusion) at the stage of the double-positive thymocyte, the unique CDR3 sequences constitute a molecular barcode identifying a single clone of the T-cell (25). Unlike the mutational heterogeneity which is constantly changing during tumour evolution by hypermutation of cancer genomes, the clonotypic heterogeneity can only be generated at the level of pre-malignant precursor T-cell, because the essential recombinases RAG1/2 that mediate V, D, J rearrangements are neither expressed in mature T-cells nor in tumour lymphocytes of MF.

We identified malignant TCRβ clonotypes by matching their frequency to TCF of the samples and noticed that epidermal samples had a higher TCRβ clonotype diversity in comparison to the dermis (median of 25 clonotypes (range 1-70) versus 11 clonotypes (range 7-49), respectively) (Fig 1A,B). However, the number of shared clonotypes between epidermis and dermis was very low, from no shared clonotypes (sample MF17), 1 shared clonotype (MF41) to a maximum of 2-5 clonotypes (MF18, MF22, MF23, MF28 and MF42) (Fig 1C). These results indicated that the pools of malignant T-cells in the epidermis and dermis are largely clonotypically unrelated that in turn suggested that they originate from separate seeding events by different T-cell clones. Indeed, we observed that in 4 of 6 samples analyzed, T-cells from the epidermal and dermal compartments individually shared between 1-8 TCRβ clonotypes with those in the circulating blood (Fig 1D), which represented a higher degree of overlap than seen for the epidermal and dermal compartments. Taken together, the epidermal and dermal compartments of the skin are likely to be seeded by different circulating malignant clones.

**Figure 1:**
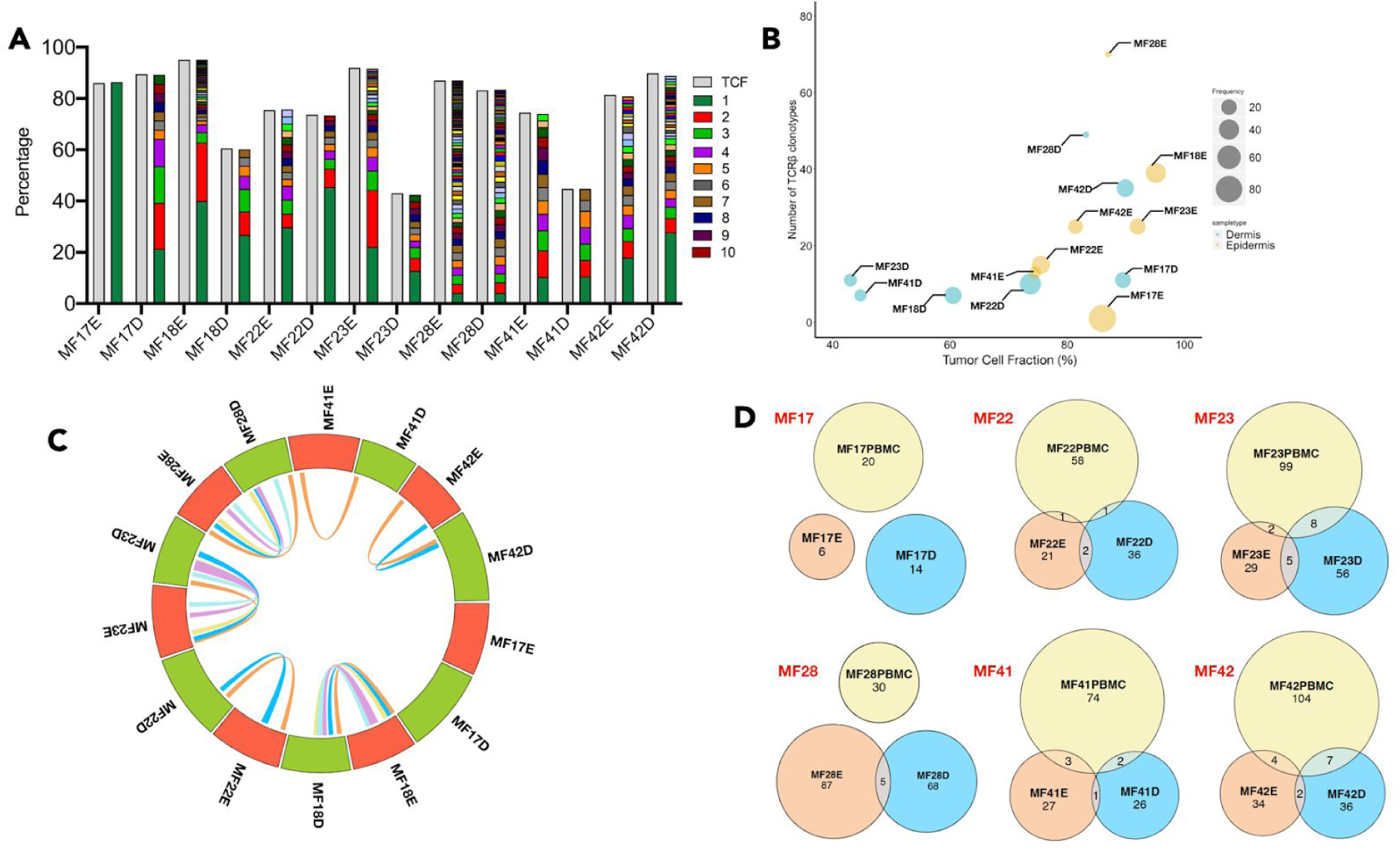
Clonotypic heterogeneity and tumour cell seeding of the skin microenvironment in MF. Percentage of Tumor cell fraction (TCF) and relative frequency of TCRβ clonotype sequences for cells isolated from different skin layer (epidermis and dermis) was calculated and plotted as a bar graph (**A**) The green and brown colour indicate the first and the 10th most frequent TCRβ clonotype in the sample. Gray colour indicates the tumour cell fraction (TCF). (**B**) Bubble plot presenting the correlation between TCF and the number of neoplastic TCRβ clonotypes in cells from epidermis and dermis of each sample. The size of the bubble is equivalent to the relative frequency of the most frequent TCRβ clonotype in the sample. (**C**) Circos plot indicates the frequency of TCRβ clonotype for cells isolated from epidermis and dermis of each sample. The connecting lines inside indicate the number of overlapping TCRβ clonotype between the two regions of the same sample. E-Epidermis; D-Dermis. (**D**) Venn diagram indicating the number of identical TCRβ clonotype between the epidermis, dermis and the circulating blood in samples MF17, MF22, MF23, MF28, MF41 and MF42.

### Mutational diversity in neoplastic T-cells in epidermis and dermis

The substantial clonotypic discordance between the epidermal and dermal compartments prompted a question regarding differences and similarities in their mutational evolution. In our previous work, we characterized 75 putative driver mutations involved in the pathogenesis and progression of MF (17). Similarities in the patterns of driver mutations between the epidermal and the dermal infiltrate would suggest parallel evolution in both compartments whereas lack of substantial overlap would indicate a neutral evolution.

We identified a median of 856 non-synonymous mutations in cells from the dermal region and 1431 non-synonymous mutations in cells from the epidermal region (Fig 2A). The majority of the mutations (48-93%) were in the Pautrier microabscess fraction and the overlap between the compartments was less than 7% across all 7 samples (supplementary **Fig 2B**). When driver genes were considered, 37 drivers were mutated in both epidermis and dermis, 13 genes (*NCOR1, ARHGEF3, ZEB1, TP53, PLCG1, RFXAP, CD58, TNFRSF1B, JAK3, MAPK1, PRKCB, MTOR* and *NF1*) were exclusively mutated in epidermis and 9 genes (*DNMT3A, TET2, SMARCB1, KDM6A, SETDB2, STAT3, NFKB2, NOTCH2* and *CARD11*) were mutated only in dermis (Fig 2C). The mutations present only in the malignant T-cells of epidermis were in the genes involved in cytoskeletal remodelling, DNA damage and immune surveillance. However, the driver mutation profile in the epidermal and dermal fractions showed non-overlapping patterns arguing against parallel evolution. Thus, the data supported the model of independent mutational evolution of neoplastic cells in different skin microenvironments.

**Figure 2:**
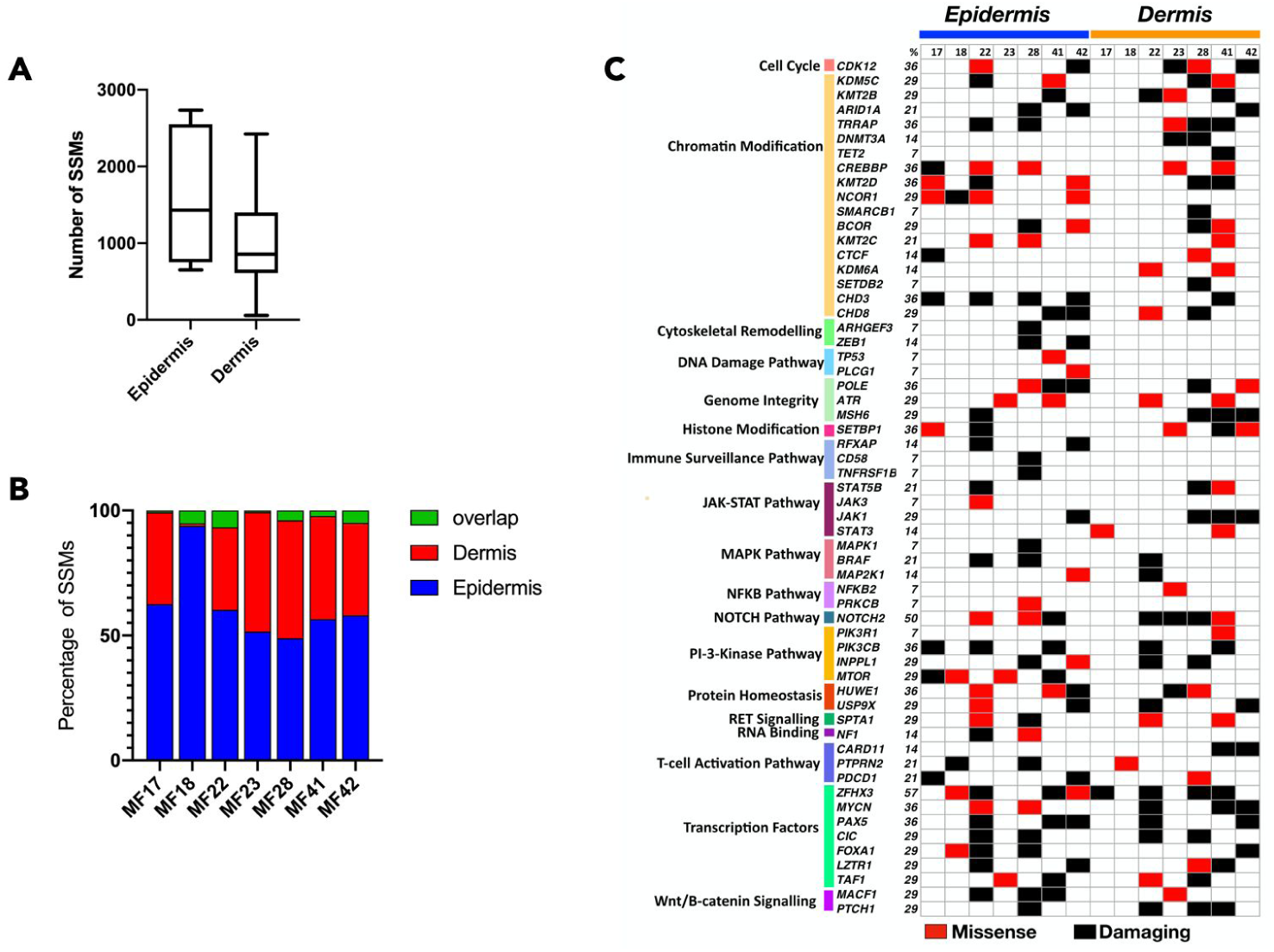
Mutational landscape of putative driver genes in anatomical layers of skin. Neoplastic T-cells isolated from epidermis and dermis were analyzed for somatic variants (SVs) in putative driver genes. Mutations in 59 genes across 18 different pathways were identified. The mutations were classified as missense or damaging. Frameshift, insertion or deletion (<6bp), stop gain or lost are classified as damaging as these mutations are likely to be deleterious.

### Phylogenetic development of tumor T-cells in skin microenvironment

To further examine the phylogenetic relationships between the subclones in the epidermal and dermal compartments we adopted the previously described bioinformatic approach based on the analysis of the mutational pattern between cancer cells (17). We found evidence of subclonal heterogeneity in all samples confirming previous findings of ITH in MF (17). A slightly higher number of subclones were found in epidermal (5-8 subclones) versus the dermal layers (4-5 subclones) (Fig 3A) reflecting the differences in clonotypic richness between those compartments (Fig 3B). In the epidermal fraction, the mutational burden was mostly in the clades whereas the dermal fraction tended to have a higher proportion of clonal (stem) mutations (Fig 3C). Thus, the number of subclones correlated with the proportion of subclonal mutations, as predicted for the neutral, branched evolution pattern (25). We also analyzed driver gene mutations in the stem and clade population for the dermis and epidermis and found that mutations in *STAT5B* and *CDK12* where only present in clades in Pautrier microabscesses whereas *NOTCH2* and *PRKCB* mutations were only in the dermal fraction, either in the stem or clades (Fig 3D).

**Figure 3:**
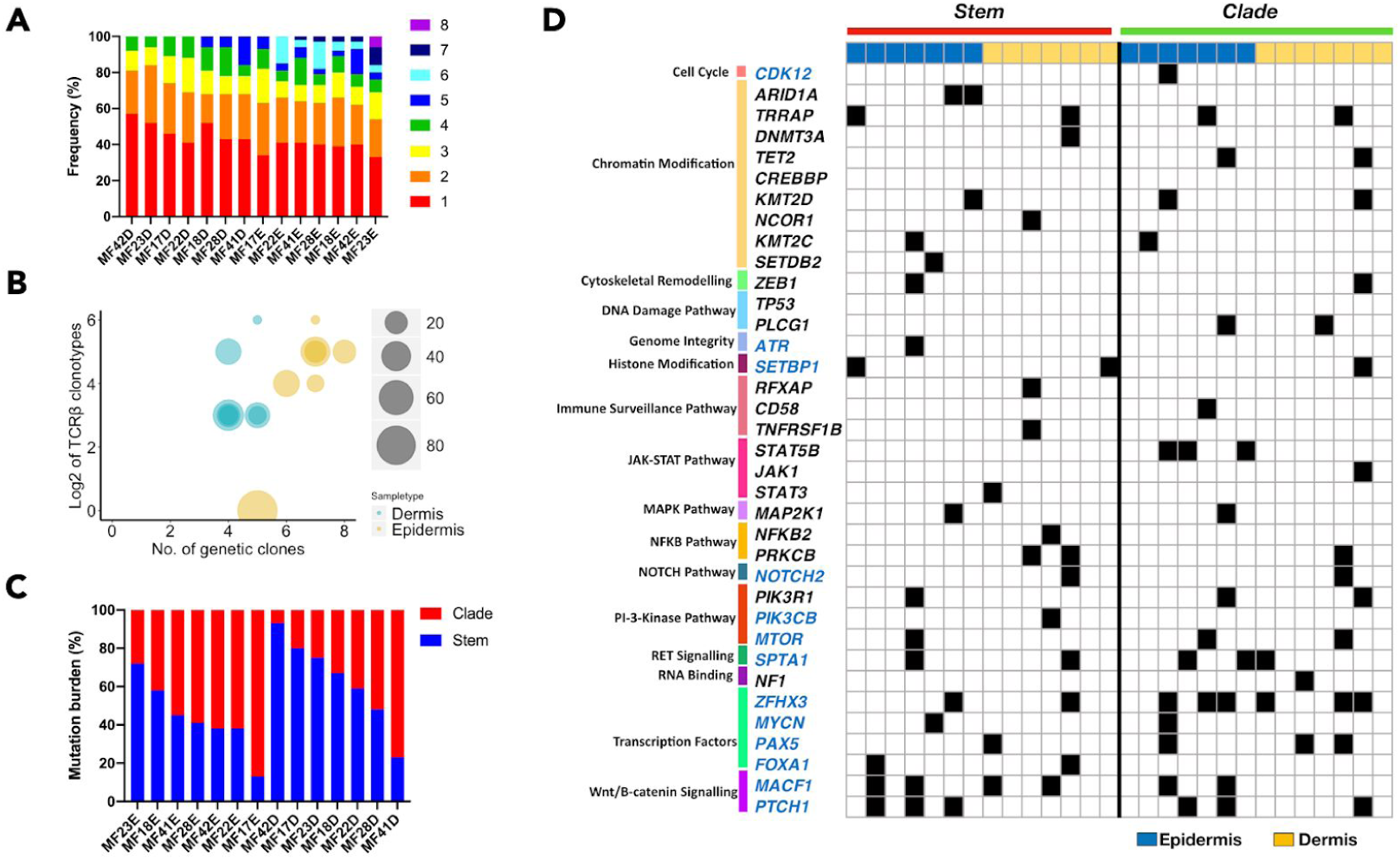
Intratumoral heterogeneity of MF in skin microenvironments. Combined data from SVs and CNA for each sample was subjected to phylogenetic analysis to identify genetic subclones. (**A**) Rainbow graph representing the number and proportion of the subclones identified in each sample. (B) Bubble plot representing the correlation between the TCRβ clonotypes and the genetic subclones. The number of TCRβ clonotypes are represented as Log2 scale. (**C**) Phylogenetic trees are composed of stem and clades (also recognized as branches). Bar graph represents the percentage of all mutations in each section (stem and clade) of the phylogenetic tree. The blue and red colour represents the mutations in stem and clades respectively (**D**) Mutational landscape of the putative driver genes in the different sections of the phylogenetic tree for two layers of skin (epidermis and dermis). Function significance of the mutations include missense, frameshift, insertions, deletions, stop gain or loss and variant in 3′ and 5′ UTR. No colour indicates absence of mutation in the sample.

To visualize how different subclones in the clades are related to each other, we reconstructed the phylogenetic trees. In one case (MF41) there was no common ancestor clone linking epidermal and dermal subclones. In other cases we detected 1-2 subclones forming the stem of the tree. All samples showed branched evolution of the subclones, with the epidermal and dermal clades clearly separated from each other (Fig 4).

**Figure 4:**
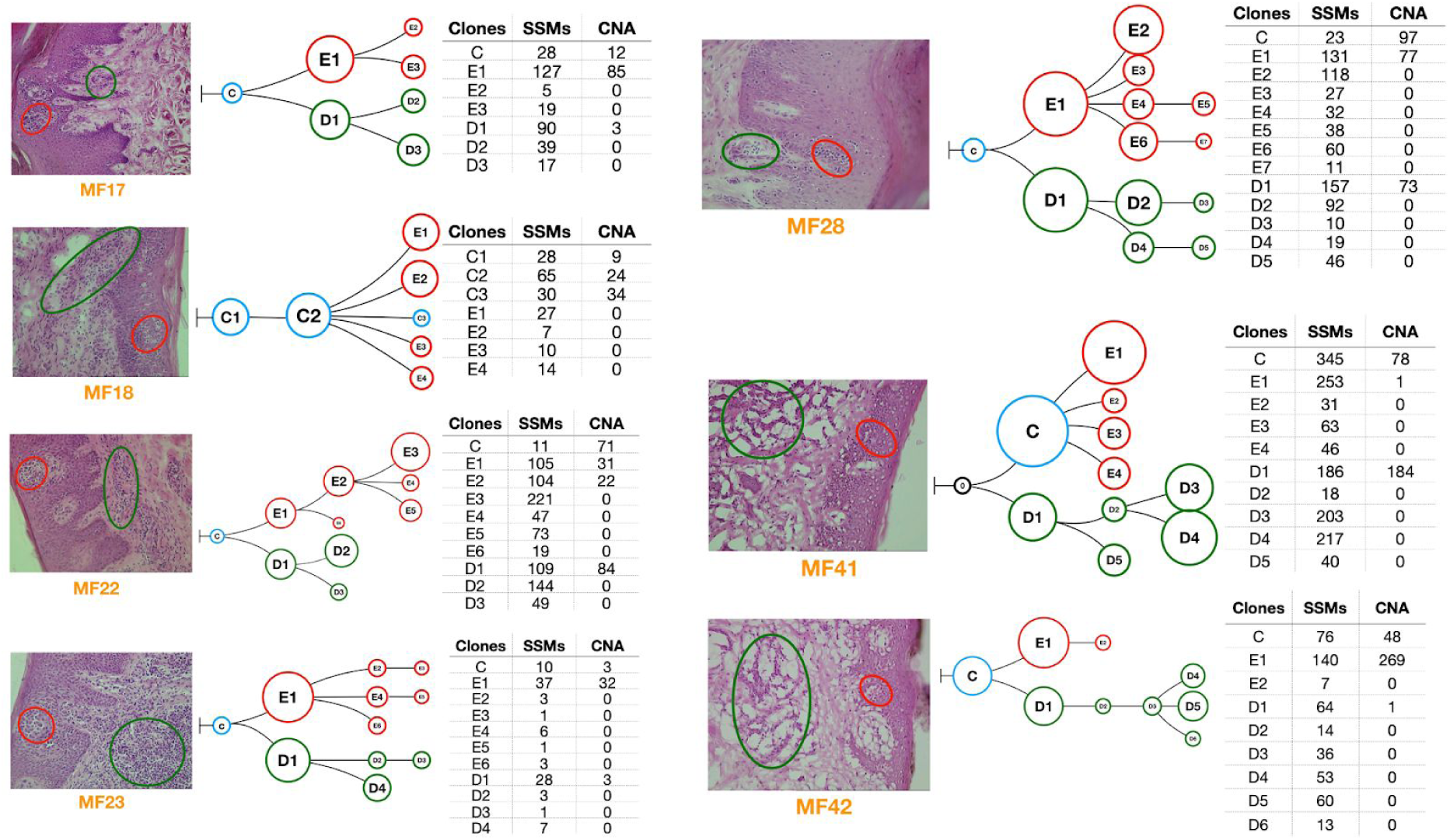
Phylogenetic analysis of the neoplastic T-cells in skin microenvironment. Genetic abnormalities (SVs and CNA) for neoplastic cells microdissected from epidermis and dermis were subjected to phylogenetic analysis. Each phylogenetic tree represents an individual patient sample. The blue circles indicate the common clone between the two skin layers. Red and green indicate the subclones in epidermis and dermis respectively. Black circles indicate absence of common ancestral clone. The tables adjacent to each figure provides the number of SVs and CNA identified in each of the subclones in the phylogenetic tree.

## Discussion

Many normal tissues comprise a system of morphologically and functionally distinctive niches that differ by their cellular composition, extracellular matrix, metabolic conditions, and accessibility to the immune system. Although tissue microenvironment has been recognized as a major factor that influences tumour cell morphology and function (27, 28), the impact of the niche on ITH and mutational evolution is poorly understood and largely limited to metastasis (29, 30).

These results help to understand how the distinct microenvironments of the skin influence the evolution of MF, a primary cutaneous, extranodal T-cell lymphoma. Our previous research showed that MF is clonotypically and genetically diverse exhibiting a high degree of ITH. The main mechanism responsible for the heterogeneity is the hematogenous seeding of skin lesions by clonotypically diverse neoplastic T-cells. The finding that the epidermal and dermal layers of the skin comprise distinct malignant clonotypes allowed us to conclude here that those compartments had been colonized by different clones of cancer cells (in this context, we define the clone as a population of malignant T-cells that are derived from the common precursor cell and exhibit the same TCRβ clonotype). Thus, the lesion of MF does not develop via a gradual infiltration of the tissue by the expanding tumour, but by independent microinvasion events in which different niches in the skin are colonized independently by various T-cell clones (Fig 5 and supplementary **Fig S1A**). Our conclusion was further confirmed by the finding that clonotypic diversity of the intraepidermal malignant cells exceeded the diversity found in the dermis. If the infiltration of the epidermis had been caused by some clones in the dermal infiltrate, the opposite phenomenon would have been found, i.e. higher number of malignant clonotypes in the dermis and a smaller number of epidermal clonotypes overlapping with the dermal clones. These findings reinforce and broaden the concept of epidermotropism in MF, which originally described the morphological impression of movement of malignant T-cells from the epidermis to the dermis. It seems that epidermotropism is a feature of early malignant T-cell clones which seed the epidermis more readily than the dermis.

**Figure 5:**
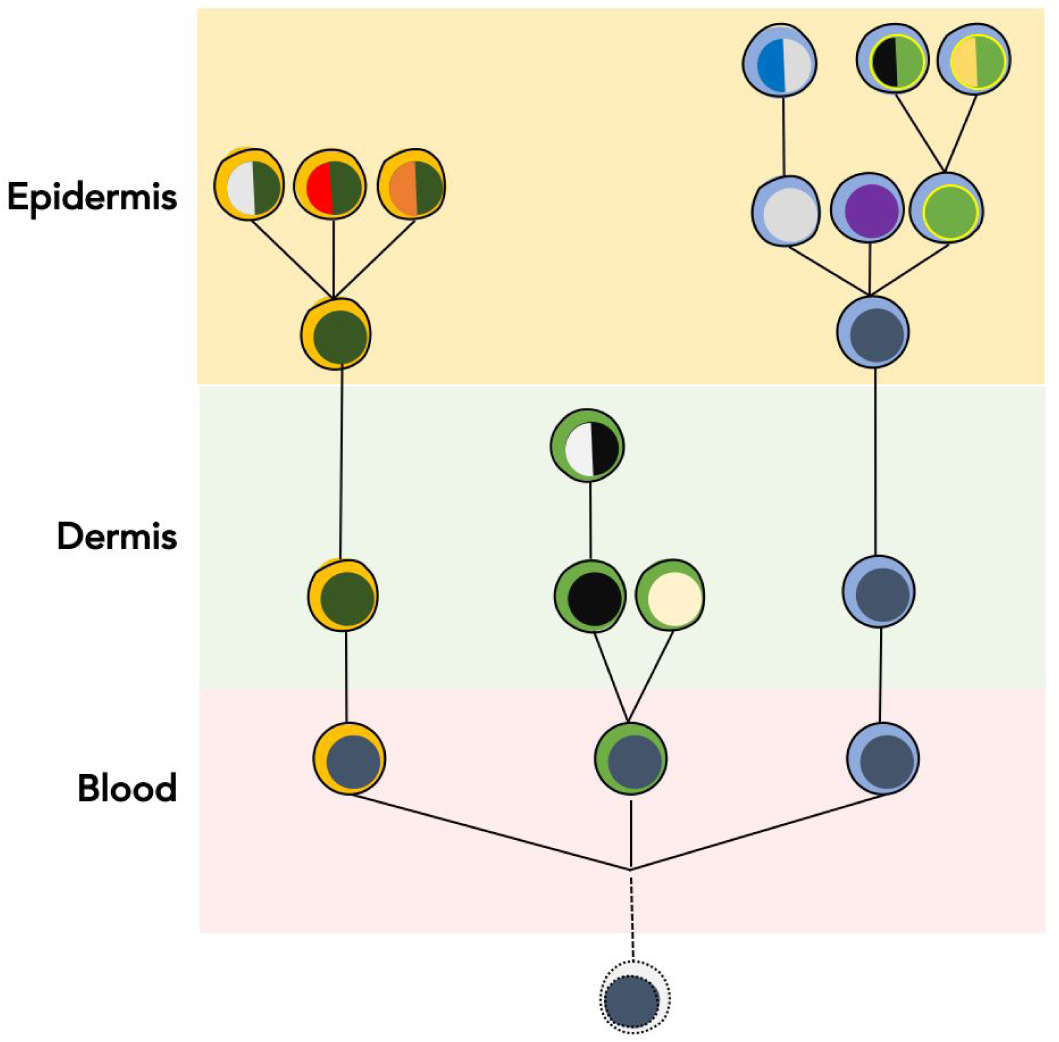
Generation of ecological heterogeneity in MF. Skin lesions of MF are initiated by circulating, clonotypically heterogeneous malignant T-cell clones (various clonotypes are highlighted by different colours of the “cytoplasm”). Upon entering the skin some clones remain in the dermis where they proliferate whereas others pass directly to the epidermis. Expanding clones accumulate mutations leading to emergence of genetically different malignant subclones (different colours of the “nucleus”). Solid lines symbolize the phylogenetic relationship between the generations of malignant cells and illustrate divergent, neutral evolution of the subclones. Based on data in this paper and our previous work (15–17, 36).

We were also able to conclude that the existence of different skin niches colonized by cancer facilitates the development of mutational subclones and augments ITH. By analyzing the phylogenetic trees of MF we found that cancer subclones seem to develop independently in each compartment via a neutral, branched evolutionary process. Similar patterns of neutral evolution have previously been found in other solid neoplasms such as the lung or colorectal cancers (26,31). The symmetric shapes of the trees suggested lack of perceptible competition between epidermal and dermal subclones. We postulate therefore that evolving subclones in different compartments do not directly compete for the niche or the nutrients and develop independently adding to the overall ITH of the lesion. Of note, lack of competition does not imply lack of interactions between different subclones. It is likely (although at this stage still hypothetical) that different MF subclones cooperate to achieve optimal tumour growth in a similar manner to what has already been shown for solid cancers (32).

The described independent evolution of cancer subclones in different tissue microenvironments is reminiscent of the well-known phenomenon of ecological speciation during the evolution of organisms, in particular, the so-called peripatric speciation (33,34). It occurs when a small fraction of the population becomes separated into a new environment. The major difference, however, is the nature of the isolation. Rather than reproductive isolation essential for speciation, different microenvironments of the tissue separate evolving subclones protecting them against direct competition. The result is a more rapid increase in ITH that what could be achieved in a homogenous environment. We would like to propose the term “ecological heterogeneity” to describe the difference in cancers subclones in various microenvironments of the tissue.

High ITH of the tumors has been correlated to unfavourable prognosis over a large range of cancers (5). Currently, data are too limited to be able to investigate the prognostic role of ITH in MF. Some indirect evidence, such as the correlation between the presence of Pautrier microabscesses with the risk of progression (35) and higher ITH in MF tumors as compared to early plaques (17) suggest that this indeed may be the case. One of the indications that ecological heterogeneity in MF may have functional significance is the non-overlapping pattern of driver mutations in epidermal and dermal neoplastic cells, which potentially would limit the efficacy of targeted therapies. However, a more detailed understanding of the differences between functionally significant signalling pathways on the level of transcriptome and the protein is needed.

## Supporting information

Independent evolution of cutaneous lymphoma subclones in different microenvironments of the skin

## Acknowledgement

We would like to acknowledge Dr. Thomas Salopek, Mrs. Rachel Doucet and the nursing staff of Edmonton Kaye Clinic for their help in sample collection. This study was supported by grants from the following sources: Canadian Dermatology Foundation (CDF RES0035718), University Hospital Foundation (University of Alberta), Bispebjerg Hospital (Copenhagen, Denmark), Danish Cancer Society (Kræftens Bekæmpelse R124-A7592 Rp12350) and an unrestricted research grant to R.G. from Department of Medicine, University of Alberta.

## Authors contribution

AI designed the experiments, analyzed the data, wrote the manuscript and submitted data to dbGAP. AI and SO performed the experiments. DH and JP performed the bioinformatic analysis. WW and GW provided input with the technical aspects of the experiments, bioinformatic pipelines, and edited the manuscript. RG designed and supervised the experiments, data analysis and edited the manuscript. All authors approved the final version of this paper.

