## Supplementary material for "Independent evolution of cutaneous lymphoma subclones in different microenvironments of the skin": Independent evolution of cutaneous lymphoma subclones in different microenvironments of the skin

### Supplementary figures and tables

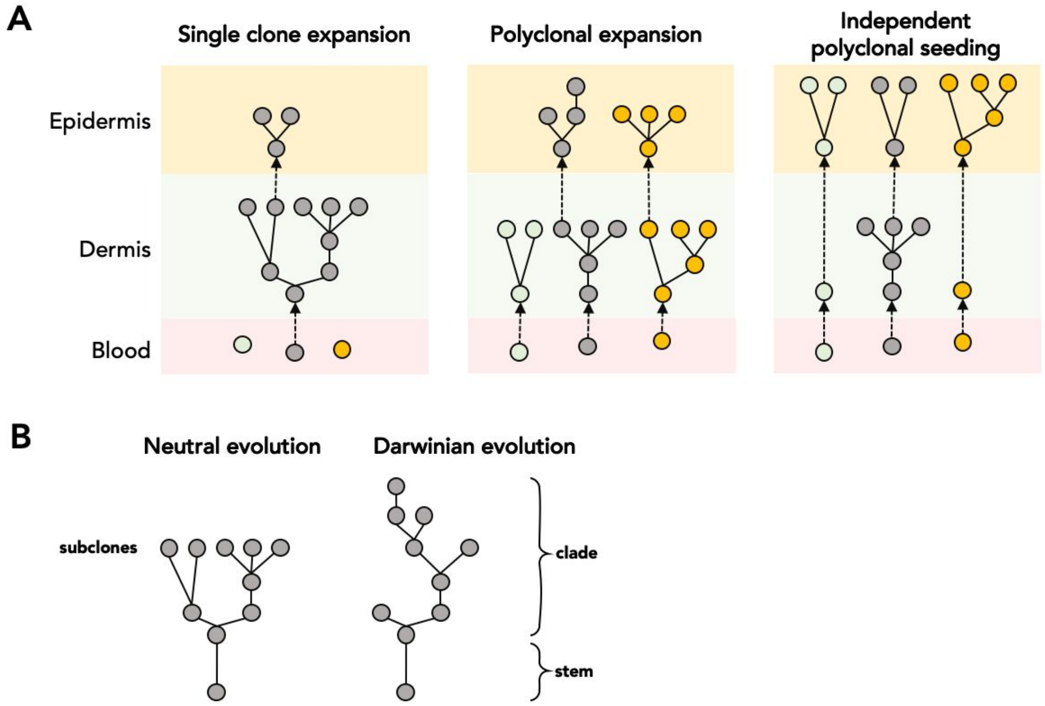


**Figure S1: Possible scenarios of tumor evolution and generating intratumoral heterogeneity in MF. (A)** Our previous work showed that lesions of MF are initiated by polyclonal circulating malignant T-cells homing to the skin where they undergo expansion and accumulation of mutations (1,2). Various clones (defined as T-cells sharing identical TCRβ clonotype) are highlighted by different colours. Expanding clones accumulate mutations and form subclones forming a phylogenetic structure. In lesions funded by a single T-cell clone (left), the entire lesion will comprise the same clonotype and the epidermal subclones will form a branch of the phylogenetic tree. If the lesion is initiated by diverse subclones (middle) that primarily proliferate in the dermis and secondarily infiltrate the epidermis the malignant cells in the epidermis and dermis would be polyclonal but epidermal malignant T-cells will form branches derived from dermal subclones. Finally, in the case of independent seeding of the dermal and epidermal niches (right), both compartments will harbour cells showing non-overlapping (or partially overlapping) clonotypes and independent patterns of mutational subclones. **(B)** Different shapes of phylogenetic tree characteristic for neutral evolution and

Darwinian evolution by natural selection. In neutral evolution, the shape of the phylogenetic tree is symmetrical (left) in contrast to Darwinian evolution where the extinction of the subclones will prune some branches (right). To avoid confusion we use the term “clone” as the group of T-cells of identical clonotype (i.e. sharing a common ancestry) rather than to mutationally identical groups of cells which we refer to as “subclones”. Clades are collections of several subclones.

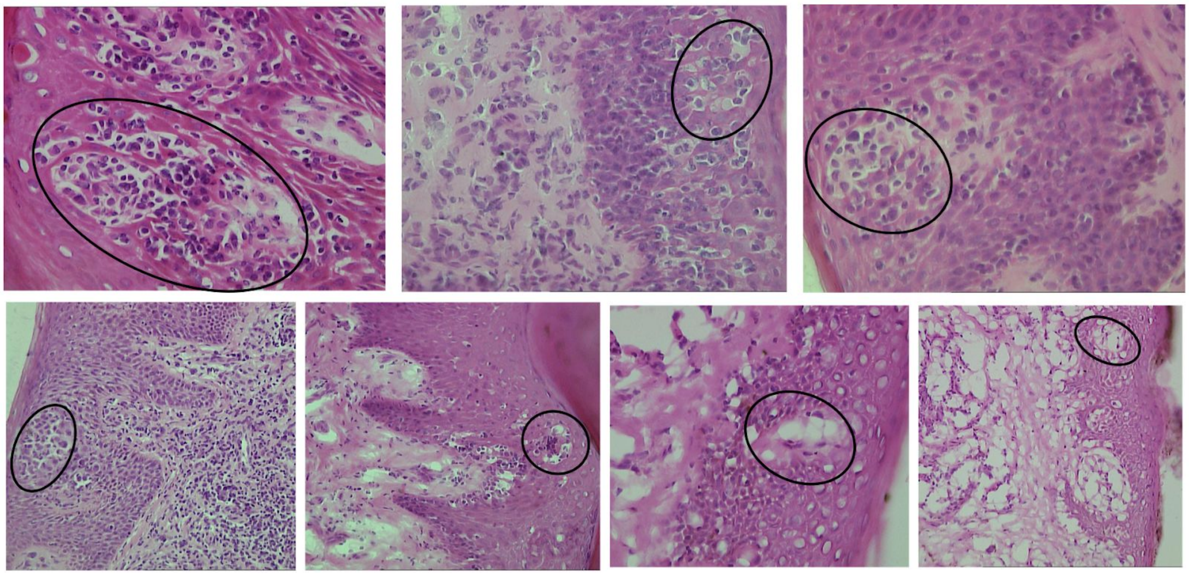

**Figure S2: Histological identification of Pautrier microabscess in MF.** Skin biopsies of MF lesions were sectioned at 10μ and stained with hematoxylin and eosin. Pautrier microabscesses are clearly discernible as clusters of atypical lymphoid cells with enlarged hyperchromatic nuclei in epidermis (black ovals). Representative images (magnification of 20x or 40x) of the H and E stained issues are presented for the 7 patients in the study.

**Supplementary Table S1:** Patient characteristics and samples included in the study

| Patient ID ( age [years], sex [M-male, F-female]) | Sample ID | Lesion type | Diagnosis and stage |
|---|---|---|---|
| MF17 (70,M) | MF17E | Plaque | Mycosis Fungoides, IB |
| | MF17D | | |
| MF18 (78, M) | MF18E | Plaque | Mycosis Fungoides, IB |
| | MF18D | | |
| MF22 (56, F) | MF22E | Plaque | Mycosis Fungoides, IA |
| | MF22D | | |
| MF23 (69, F) | MF23E | Plaque | Mycosis Fungoides, IA |
| | MF23D | | |
| MF28 (65, M) | MF28E | Plaque | Mycosis Fungoides, IB |
| | MF28D | | |
| MF41(77, F) | MF41E | Plaque | Mycosis Fungoides, IIB |
| | MF41D | | |
| MF42 (82, M) | MF42E | Plaque | Mycosis Fungoides, IIIB |
| | MF42D | | |

2. Iyer A, Hennessey D, O'Keefe S, Patterson J, Wang W, Wong GK-S, et al. Branched evolution and genomic intratumor heterogeneity in the pathogenesis of cutaneous T-cell lymphoma [Internet]. Available from: http://dx.doi.org/10.1101/804351
